# Fibroblastic cells are a site of mouse cytomegalovirus *in vivo* lytic replication and latent persistence oppositely regulated by *Stat1*

**DOI:** 10.1101/2022.06.29.498076

**Authors:** Katarzyna M. Sitnik, Fran Krstanović, Natascha Gödecke, Ulfert Rand, Tobias Kubsch, Henrike Maaß, Yeonsu Kim, Ilija Brizić, Luka Čičin-Šain

**Affiliations:** Department of Viral Immunology, Helmholtz Centre for Infection Research, 38124 Braunschweig, Germany; Institute of Animal Breeding and Genetics, University of Veterinary Medicine Vienna, 1210 Vienna, Austria; Center for Proteomics, Faculty of Medicine, University of Rijeka, 51000 Rijeka, Croatia; Centre for Individualized Infection Medicine, a joint venture of HZI and MHH, 30625 Hannover, Germany; German Centre for Infection Research (DZIF), Hannover-Braunschweig site, 38124 Braunschweig, Germany

**Author notes:** Senior author (Katarzyna M. Sitnik, Luka Čičin-Šain). Corresponding author, Correspondence: Luka Čičin-Šain, (Lead contact) and Katarzyna Sitnik.

## Abstract

To date, no herpesvirus has been shown to latently persist in fibroblastic cells. Here, we demonstrate that mouse CMV (MCMV), a β-herpesvirus, persists for the long term and across organs in PDGFRα^+^ fibroblastic cells, with similar or higher genome loads than in the previously known sites of MCMV latency. Whereas MCMV gene transcription in PDGFRα^+^ fibroblastic cells was almost completely silenced at 5 months post-infection, these cells gave rise to reactivated virus *ex vivo*, arguing that they supported latent MCMV infection. Notably, PDGFRα^+^ fibroblastic cells also supported productive virus replication during primary MCMV infection. Mechanistically, *Stat1*-deficiency resulted in increased lytic but abolished latent infection of fibroblastic cells *in vivo*. In sum, fibroblastic cells have a dual role as a site of lytic MCMV replication and a reservoir of latent MCMV *in vivo* and *Stat1* is critically involved in the regulation of MCMV latency.

## Introduction

Mouse CMV (MCMV) belongs to the genus *Muromegalovirus* in the subfamily *Betaherpesvirinae*. It is most closely related to the primate-infecting viruses of the genus *Cytomegalovirus*, comprising human CMV (HCMV). Together, CMVs are characterized by large DNA genomes (>200 kb), induction of cytomegaly in infected cells and strict species tropism [1]. As for all herpesviruses, the life cycle of CMVs comprises both lytic (productive) and latent (non-productive) infection, the latter underpinning their ability to persist for life in their respective hosts. During lytic infection the CMV genome is transcribed to express high levels of immediate–early (IE), early (E) and late (L) viral gene products, added in a temporal sequence, that carry the virus through the phases of genome replication and virion assembly, culminating in the release of infectious virions and cell lysis. By contrast, during latent infection CMV genomic DNA persists in the cell nucleus but the viral progeny is not produced. Viral gene transcription in latency is considerably silenced and the major CMV IE promoter is associated with histones bearing repressive post-translational marks [2, 3]. Under appropriate conditions, the virus productive cycle is reactivated, leading to the genesis of new infectious progeny. A formal criterium of latency is the appearance of infectious virus upon, but not prior to, co-cultivation of the population of interest with cells permissive to lytic infection. *In vivo*, lytic CMV infection prevails for some time following virus inoculation, enabling viral amplification and spread, but in immunocompetent hosts it is eventually extinguished, after which a life-long phase of *in vivo* latency ensues when only latent virus can be detected [4].

Herpesviruses are recognized for using distinct cell types as sites of lytic replication and latent persistence [5]. In line with this, myeloid progenitors and monocytes are well-characterised sites of HCMV latency [6]. Latent infection of these cell types by MCMV is less clear, due to conflicting results in studies [7, 8] and remains a controversial topic. On the other hand, and somewhat curiously for a herpesvirus, latent MCMV was found to reside during *in vivo* latency in liver endothelial cells [8] and peritoneal macrophages [9, 10], which represent cell types that are permissive to lytic MCMV infection [9, 11]. Fibroblasts are also well-known to support lytic MCMV replication and are commonly used to grow virus stocks [12]. Interestingly, experiments of Mercer *et al*. [13] and Pomeroy *et al*. [14] performed more than 30 years ago, demonstrated enrichment for MCMV genomes and virus reactivation from stroma-enriched spleen fragments, while essentially excluding the contribution of hematopoietic cells. The authors proposed that splenic fibroblasts were involved but could not prove it due to the lack of available methods for splenic fibroblast purification. Notably, the notion about latency in fibroblastic cells has since not been investigated for any CMV or other herpesvirus, and remains unconfirmed.

Like other viral infections, MCMV is crucially controlled by the host’s response to interferons (IFNs) [15], which signal through signal transducer and activator of transcription 1 (STAT1) [16]. Mice deficient for *Stat1* or otherwise rendered unresponsive to IFNs succumb within a few days to infection with only a few hundred plaque forming units (PFU) of MCMV [17, 18]. Besides orchestrating antiviral immunity through effects on the functions of the cells of the immune system, IFNs directly inhibit productive MCMV growth by elevating cell-intrinsic antiviral activity [15, 19, 20]. Notably, IFN treatment inhibits the activity of MCMV major IE promoter, which controls the initiation of the lytic cycle [20]. We have shown that MCMV infection in permissive cells infected at low multiplicity of infection (MOI) in the presence of IFN is transiently repressed until IFN is removed. This reversible state satisfied the formal definition of latency, where MCMV genomes are maintained in absence of replicating virus, but the virus has the capacity to reactivate [20]. However, it remained unclear if similar phenomena may be observed in a long-term *in vivo* model of CMV latency.

In this study, using state-of-the-art methods for the purification of tissue-resident subsets from mouse organs, we comprehensively mapped the cellular sites of MCMV genome persistence during latency *in vivo*. Our study unravels that MCMV commonly persists in PDGFRα^+^ fibroblastic cells for the long term, in a state that fulfils all the formal criteria of latent infection. We further show that PDGFRα^+^ fibroblastic cells are simultaneously permissive to lytic MCMV infection and have a role in viral reproduction during the acute phase of *in vivo* infection. Finally, we identify a physiological role of *Stat1* in promoting latent persistence of MCMV in fibroblastic cells *in vivo* whilst concurrently inhibiting their lytic infection with the virus.

## Results

### PDGFRα^+^ fibroblastic cells carry MCMV genomes during latent MCMV infection *in vivo*

To address whether fibroblastic cells might be a site of MCMV latent persistence *in vivo*, we initially performed a comprehensive characterisation of the distribution of MCMV genomes in tissue-resident cells across mouse organs during latency *in vivo*. Specifically, we determined the number of viral genome copies per 100,000 cells (henceforth referred to as MCMV genome load) in FACS-sorted fibroblastic-, endothelial-, epithelial- and tissue-resident macrophage populations from the liver, salivary glands (SG), spleen, visceral adipose tissue (VAT) and lungs of mice latently infected via intraperitoneal route (Fig. 1a). Intraperitoneal administration of MCMV is the most widely used MCMV infection model that results in infection of multiple organs due to virus systemic spread in the acute phase of infection [21, 22]. Populations of tissue-resident macrophages (Mϕ), such as F4/80^+^CD11c^+^ SG Mϕ, lung alveolar Mϕ, liver Kupfer cells, spleen red pulp Mϕ and VAT Mϕ were identified using organ-specific gating strategies [23–25]. Endothelial cells (EC) were identified as CD45^−^ PDGFRα^−^ CD31^+^ cells and were shown to co-express CD146 in the liver (Supplementary Fig. 1b). Of note, EC from VAT were not analysed as they could not be isolated in sufficient numbers. PDGFRα^+^ fibroblastic cells (PDGFRα^+^ FC) were identified as CD45^−^CD31^−^PDGFRα^+^ cells in VAT [26], CD45^−^CD31^−^EpCAM^−^ PDGFRα^+^ cells in the lungs [27] and SG [28]; and as CD45^−^CD31^−^ITGB1^+^PDGFRα^+^ cells in the spleen [29]. In the liver we focused on a defined CD45^−^CD31^−^EpCAM^−^PDGFRα^+^ FC subset, vitamin A^+^ liver stellate cells [30, 31]. We chose to focus on FC that express PDGFRα since they are relatively well-defined across mouse organs [32]. For detailed phenotypes and gating schemes for all cell populations mentioned above see Fig. 1 and Supplementary Fig. 1a. To quantify MCMV genome load, genomic DNA isolated from sorted cell subsets was used as input in a quantitative PCR assay with primers specific to viral gB gene and to mouse *Pthrp* gene. Absolute quantification was performed using serial dilutions of a plasmid carrying both gB and *Pthrp* sequences, according to a previously described method [33], with a quantitation limit of 10 copies of the plasmid standard in a single qPCR reaction (Supplementary Fig. 1c). Strikingly, PDGFRα^+^ FC carried the highest viral genome load across multiple tissues, in the liver (ca 2-fold), in SG (ca 10-fold) and most substantially, in the spleen (ca 100-fold) (Fig. 1b-i). Only in the lungs were MCMV genomes absent from PDGFRα^+^ FC (Fig. 1j-k). EC carried MCMV genomes in liver and lungs, which is consistent with previous reports [8, 34], but their MCMV genome content appeared lower in the spleen and was barely detectable in the SG (Fig. 1b-k). MCMV genomes were detected in VAT Mϕ at levels similar to PDGFRα^+^ FC from the same site, but were absent or barely detectable in Mϕ from other organs (Fig. 1b-k). Next, to compare the viral genome load in PDGFRα^+^ FC to that of circulating myeloid cells, we analysed monocytes isolated from bone marrow (BM) or the spleen of latently infected mice. We detected less than 10 copies of MCMV genomes, which is 10-100-fold lower than carried by PDGFRα^+^ FC in all organs tested except the lungs. Importantly, MCMV genomes persisted in PDGFRα^+^ FC even 1 year after infection, as shown for PDGFRα^+^ FC from the spleen, lymph nodes (LNs) [35] and VAT (Fig. 1m-o). Together, these results reveal that MCMV genomes are commonly present in PDGFRα^+^ FC of multiple organs during MCMV latency *in vivo*.

**Figure 1.**
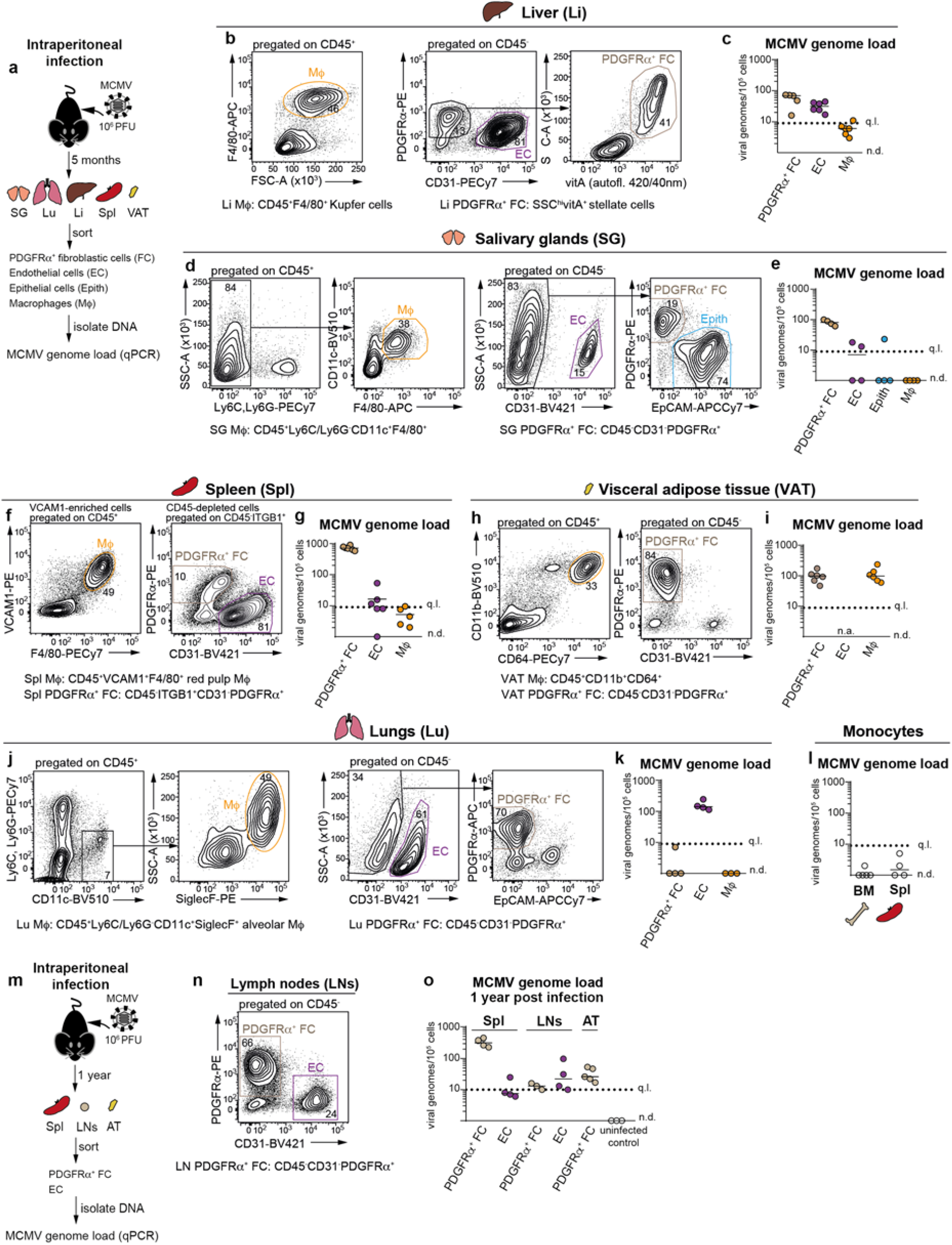
PDGFRα^+^ fibroblastic cells carry MCMV genomes during latent MCMV infection *in vivo*. Analysis of MCMV genome load in indicated cell subsets purified from mice latently infected with MCMV administered intraperitoneally **a-l**, 5 months or **m-o**, 1 year prior. **a**, Experimental set up for b-l. SG, submandibular salivary glands; Li, liver; Spl, spleen; VAT, visceral adipose tissue; Lu, lungs. BM, bone marrow. **m**, Experimental set up for n-o. Lymph nodes (LNs) represent pooled inguinal, brachial, axillary, cervical and mesenteric LNs. **b, d, f, h, j, n** Gating strategy for sorting of the indicated cell subsets. Numbers are percentage of cells in the indicated gates. **c, e, g, i, k, l, o** Number of virus genomes per 100,000 cells. Horizontal lines depict median from n = 3-6 biological replicates (depicted as symbols). Biological replicates represent cells sorted from pooled preparations from **c, k**, 1-2 mice; **e, i, l**, 2 mice; or **g, o**, 4 mice and are from 2 independent experiments per organ. Sorting was done either from 1 organ (VAT) or simultaneously from 2 organs (SG and Lu; Li and BM; SPL and LNs). q.l., quantification limit; n.d., not detected; n.a., not analysed

### The route of infection determines the load of MCMV genomes in PDGFRα^+^ FC of latently infected lungs

Our finding that following intraperitoneal infection MCMV genomes persist in PDGFRα^+^ FC in all organs except the lungs was intriguing. We previously compared mucosal routes of infection, such as intranasal or intragastric, with intraperitoneal infection and observed that intranasal infection is the more efficient route of MCMV infection for the lungs [36]. Nevertheless, intranasal infection results in a more restricted pattern of virus spread than the intraperitoneal infection route, involving only the lungs and SG, which was in line with a previous report [37]. Notably, the cellular sites of MCMV latency after intranasal infection, which is thought to better model the natural MCMV infection [38], are yet to be investigated. Therefore, we next determined MCMV genome load in PDGFRα^+^ FC, EC as well as epithelial cells and Mϕ from the lungs and SG of mice latently infected via intranasal route (Fig. 2a).

**Figure 2.**
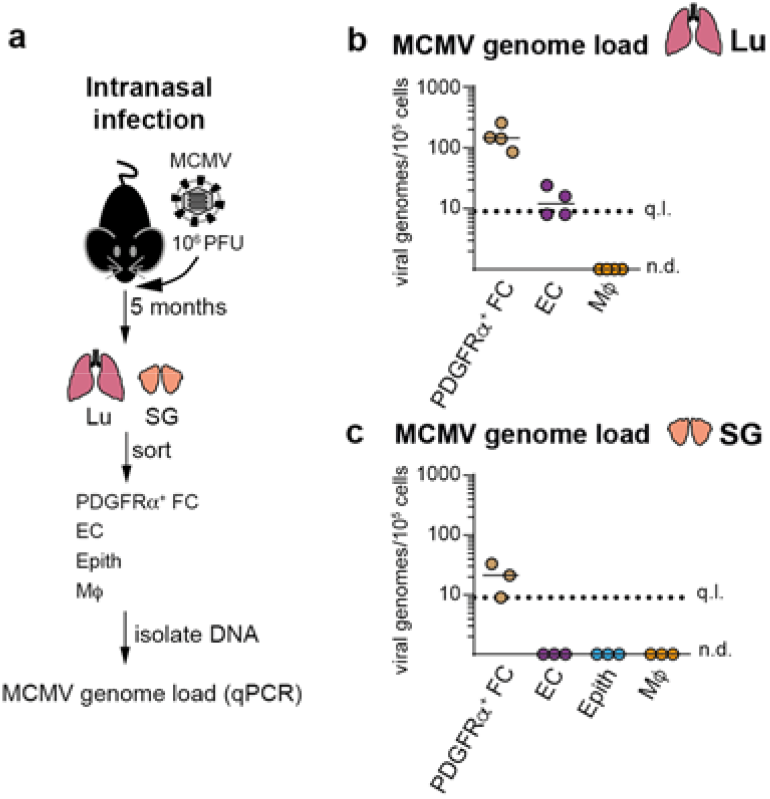
The route of infection determines the MCMV genome content of PDGFRα^+^ FC in latently infected lungs. Analysis of MCMV genome load in indicated cell subsets purified from mice latently infected with MCMV administered intranasally 5 months prior. **a** Experimental set up. **b, c** Number of virus genomes per 100,000 cells. Horizontal lines depict median from n = 3-4 biological replicates (depicted as symbols). SG and Lu subsets were simultaneously obtained from 1-2 mice per replicate. q.l., quantification limit; n.d., not detected.

Notably, PDGFRα^+^ FC carried the highest MCMV genome load of all tested subsets in both lungs and SG of mice infected intranasally (Fig. 2b-c). Thus, distinct routes of infection result in differences in MCMV genome distribution in PDGFRα^+^ FC in the lungs. Whether this is due to putative differences in the exposure/accessibility of PDGFRα^+^ FC to the virus during acute infection and/or putative differential effects on virus persistence in PDGFRα^+^ FC remains to be clarified in future studies.

### Differences in MCMV genome load between PDGFRα^+^ and PDGFRα^−^ FC during latency *in vivo*

Next, we asked whether other populations of FC, which do not express PDGFRα, carried similar MCMV genome load in latently infected mice as PDGFRα^+^ FC. To this end, we determined MCMV genome load in distinct subsets of FC from the lungs of mice latently infected either intraperitoneally or intranasally (Fig. 3a). The CD45^−^CD31^−^EpCAM^−^ cell fraction from the lungs consisted of two discrete compartments, a major subset of PDGFRα^+^ FC, which is known to comprise alveolar, peribronchial and adventitial fibroblasts [27], and a minor FC population that lacked PDGFRα but expressed PDGFRβ, hereafter termed PDGFRα^−^PDGFRβ^+^ FC (Fig. 3b). The majority of PDGFRα^−^PDGFRβ^+^ FC expressed CD146 (Supplementary Fig. 1d), fitting with the universal phenotype of mural cells [39].

**Figure 3.**
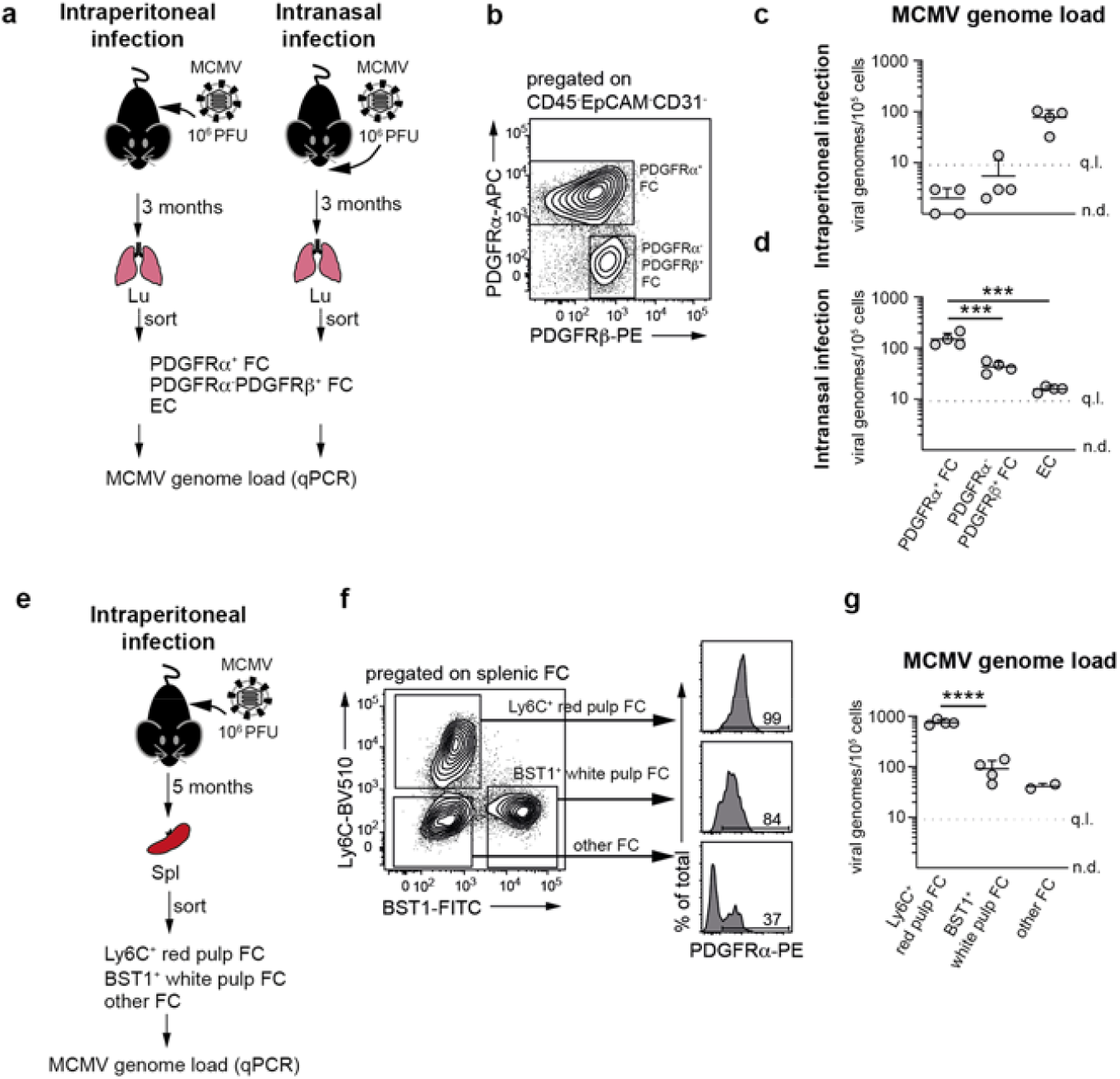
Differences in MCMV genome load between PDGFRα^+^ and PDGFRα^−^ FC during latency *in vivo*. Analysis of MCMV genome load in indicated FC subsets purified from the lungs of mice latently infected with MCMV administered intraperitoneally or intranasally (**a-d**), or the spleen of mice latently infected with MCMV administered intraperitoneally (**e-g**). **a** Experimental set up for b-d. **e** Experimental set up for f-g. **b, f** Gating strategy for sorting of the indicated cell subsets. **c, d, g** Number of virus genomes per 100,000 cells. **c, d** Horizontal line depicts the arithmetic mean ± SD from n = 4 biological replicates (depicted as symbols) pooled from 2 independent experiments. Biological replicates represent cells sorted from pooled preparations from 2 mice. ***, p-value < 0.001, one-way ANOVA with Dunnett’s multiple comparisons test **g** Horizontal line depicts the arithmetic mean ± SD from n = 2-4 biological replicates (depicted as symbols) pooled from 2 independent experiments. Biological replicates represent cells sorted from pooled preparations from 4 mice. ****, p-value < 0.0001, unpaired t test. q.l., quantification limit; n.d., not detected.

Following intraperitoneal infection, which results in the preferential distribution of MCMV genomes to EC, both PDGFRα^+^ FC and PDGFRα^−^PDGFRβ^+^ FC were a minor source of MCMV genomes with the load of less than 10 viral genome copies per 100,000 cells (Fig. 3c). Following intranasal infection, which results in the preferential distribution of MCMV genomes to PDGFRα^+^ FC, viral genomes were robustly detected in both FC subsets but the genome load in PDGFRα^−^PDGFRβ^+^ FC was ca 3-fold lower compared to PDGFRα^+^ FC (Fig. 3d). We also mapped the distribution of MCMV genomes during latency in the subpopulations of FC from the spleens of mice infected intraperitoneally (Fig. 3e). We fractionated CD45^−^CD31^−^ITGB1^+^ splenic FC [29] into Ly6C^+^ red pulp FC, BST-1^+^ white pulp FC, and BST-1^−^Ly6C^−^ FC (hereafter termed, other FC), which are mainly composed of CD146^+^ mural cells [29]. As shown in Fig. 3f, Ly6C^+^ red pulp FC uniformly expressed high levels of PDGFRα, BST-1^+^ white pulp FC contained a minor fraction of PDGFRα^−^ cells while other FC consisted mainly of PDGFRα^−^ cells. Notably, MCMV genome load was nearly 10-fold higher in Ly6C^+^ red pulp FC compared to the other two FC subpopulations (Fig. 3g). In sum, MCMV genome load is higher in the PDGFRα-positive fraction of FC in the lungs and spleen than in other FC subsets.

### Silenced and stochastic viral gene transcription in PDGFRα^+^ FC from latently infected mice

Next, we investigated if PDGFRα^+^ FC that carried viral genomes during MCMV latency *in vivo* fulfilled the formal criteria of latent infection. We first asked whether viral gene expression in PDGFRα^+^ FC from latently infected mice was consistent with a putative ongoing lytic viral cycle. To this aim, we measured the expression of viral transcripts from the immediate-early (*ie1*/*ie3*), early (*M38*) or late (*M48*/*SCP*) phase of the lytic infection in RNA isolated from PDGFRα^+^ FC from latently infected mice by RT-qPCR (Table 1). We analysed PDGFRα^+^ FC from various organs and conditions in which they were shown to carry MCMV genomes (see Fig. 1 and 2), such as the spleen, VAT, SG and liver of mice infected intraperitoneally, and from PDGFRα^+^ FC from the lungs of mice infected intranasally.

**Table 1.**
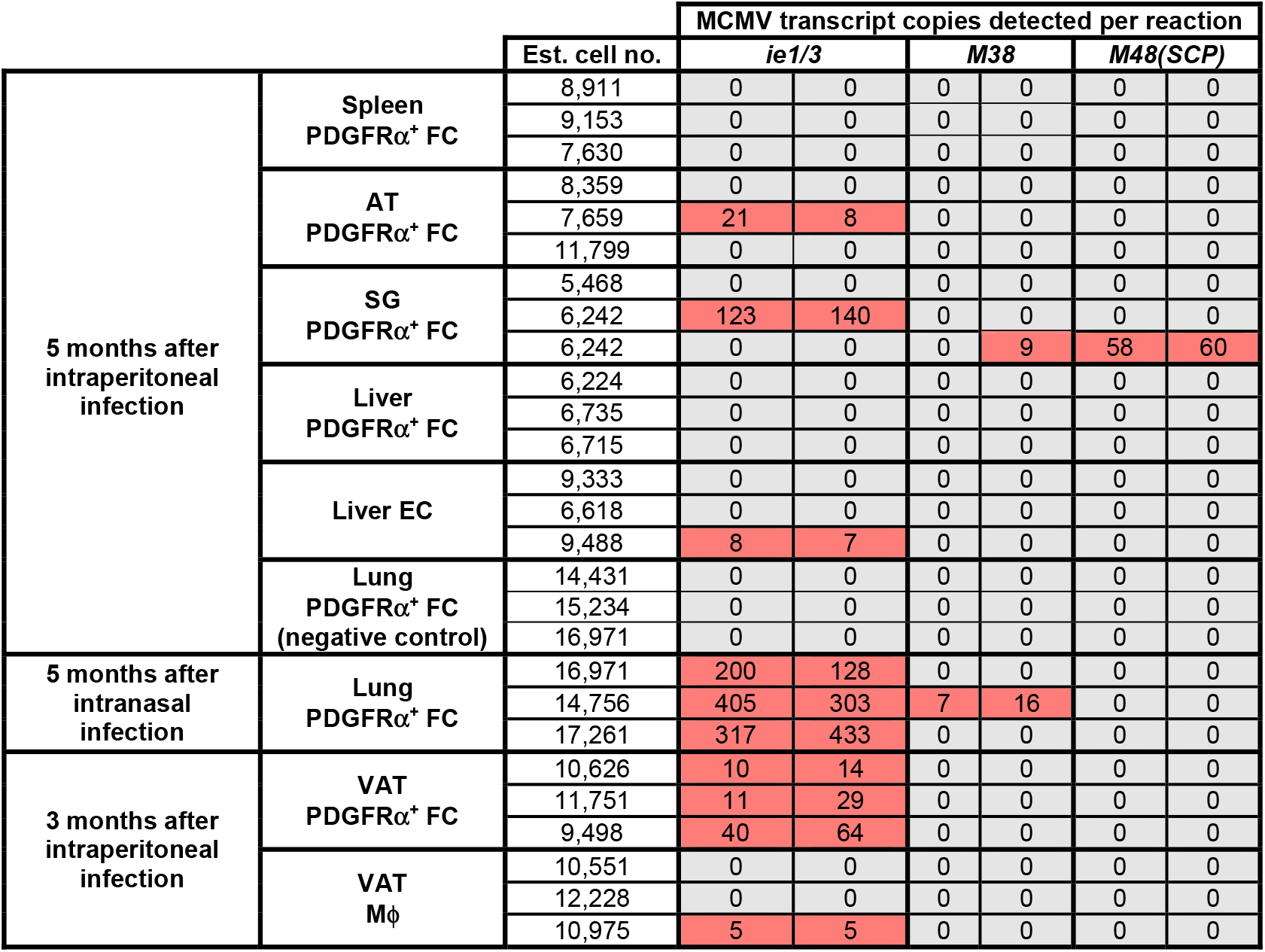
RT-qPCR analysis of the expression of MCMV genes in the indicated subsets obtained from mice latently infected with MCMV via the indicated route and for the indicated amount of time. Each row represents cDNA from a different biological replicate analysed in 2 reactions (technical duplicates) per MCMV gene. Biological replicates represent cells sorted from pooled preparations from 1-2 (Lu, Li, SG, VAT) or 4 (Spl) mice per replicate.

PDGFRα^+^ FC from the lungs of mice infected intraperitoneally, which are devoid of MCMV genomes (see Fig. 2), were used as a negative control. Liver EC and VAT Mϕ from mice infected via intraperitoneal route were also included. Following reverse transcription, copies of cDNA representing each transcript were quantified using serial dilutions of MCMV-BAC [33], with a quantitation limit of 10 copies of the BAC standard in a single qPCR reaction (Supplementary Fig. 2a). Although we used primer pairs that did not distinguish between viral DNA and RNA, our assay specifically detected viral RNA, since no positive signals were obtained when the same RNA samples were processed with omission of reverse-transcriptase (negative data not shown). To ascertain that comparable amounts of cDNA were used, we standardized our readouts on mouse *Gapdh* expression in all samples. The number of cells corresponding to the cDNA input used per qPCR reaction was estimated for each sample and was in the range of 5,000-15,000 cells (Table 1). Overall, expression of viral mRNAs was detected only sporadically and as no more than a few hundred of copies (Table 1). Specifically, *ie1*/*ie3* expression was detected in 8, *M38* in 2 and *M48/SCP* in 1 out of 18 tested PDGFRα^+^ FC samples, but none of the samples expressed all three viral genes (Table 1). This transcriptional profile is inconsistent with lytic infection while it fits with the pattern of viral transcription in latently infected lungs described recently by Griessl *et al*., which was found to be intermittent and stochastic [34]. *ie1*/*ie3* expression was detected in 1 out 3 tested liver EC samples and in 1 out 3 tested VAT Mϕ samples, indicating that viral transcription is also silenced in EC and Mϕ carrying MCMV genomes during latency *in vivo*. In sum, the profile of viral gene expression in PDGFRα^+^ FC was at odds with persistent lytic infection.

### PDGFRα^+^ FC carry latent MCMV *in vivo*

To test for functional criteria of latency, we next determined if MCMV can reactivate from PDGFRα^+^ FC purified from latently infected mice. To this aim, we devised an *ex vivo* reactivation assay, in which PDGFRα^+^ FC isolated from the lungs of mice latently infected via intranasal route were seeded in half-area 96-well plates at the density of 100,000 cells per well (resulting in the deposition of ~100 MCMV genomes per well) (Fig. 4a). Upon 7 days of *ex vivo* incubation, viral reactivation was assessed in individual wells using three criteria: i) visual inspection of plaque formation, ii) IE1 immunofluorescence as well as iii) detection of infectious virus in culture supernatant by plaque assay on monolayer of mouse embryonic fibroblasts (MEFs). Notably, infectious virus emerged in 30-50 % of wells from 4 out of 4 samples of lung PDGFRα^+^ FC (Fig. 4b-c). As expected, virus did not grow out when lung PDGFRα^+^ FC isolated from mice infected intraperitoneally were used, as they were shown to be devoid of MCMV genomes (Fig. 4d). Further importantly, no virus outgrowth was detected when lysates prepared from lung PDGFRα^+^ FC isolated from mice infected intranasally were incubated on a monolayer of MEFs for the same amount of time (Fig. 4e-f). The results showing that virus grew out only from cells but not from cell lysates (even though the assay was able to detect a single PFU of MCMV), argues that PDGFRα^+^ FC carried latent MCMV which reactivated upon *ex vivo* cultivation. To confirm this using PDGFRα^+^ FC from another tissue, but also to study the kinetic of reactivation events, virus reactivation was also monitored in PDGFRα^+^ FC isolated from the VAT of mice latently infected via intraperitoneal route. Mice were administered with reporter MCMV expressing major IE promoter-driven GFP (MCMV^IE-GFP^) [29], which resulted in the same latent viral load in VAT PDGFRα^+^ FC (Supplementary Fig. 2b) while it enabled monitoring of the kinetic of re-initiation of lytic gene transcription by life cell imaging (Fig. 4g). By 7 days of *in vitro* incubation, the virus grew out in ~10% of wells from 9 out of 10 samples (Fig. 4h-i). Virus outgrowth was specifically associated with the presence of PDGFRα^+^ FC, since cultivation of VAT cell preparations that had been depleted of PDGFRα^+^ FC did not yield any virus (Fig. 4j). It should be noted that this result does not provide evidence for the lack of reactivation from VAT Mϕ or EC since the culture conditions did not support their survival (data not shown). As shown for lung PDGFRα^+^ FC, virus was recovered only from live PDGFRα^+^ FC from VAT and not from cell lysates (Fig. 4k-l). Thus, the virus could reactivate from PDGFRα^+^ FC from both lungs and VAT, with frequency correlating with the level of IE gene transcription in these cells, which was lower in VAT than in the lungs (see Table 1). Based on the emergence of initial GFP^+^ cells, re-initiation of lytic gene transcription occurred either 1.5 - 2 days after seeding (50% of reactivations) or no sooner than 4 - 5 days after seeding (40% of reactivations) (Fig. 4i). Thus, reactivation events appeared to follow a biphasic kinetic. Together, these results indicate that MCMV latently persists in PDGFRα^+^ FC *in vivo*.

**Figure 4.**
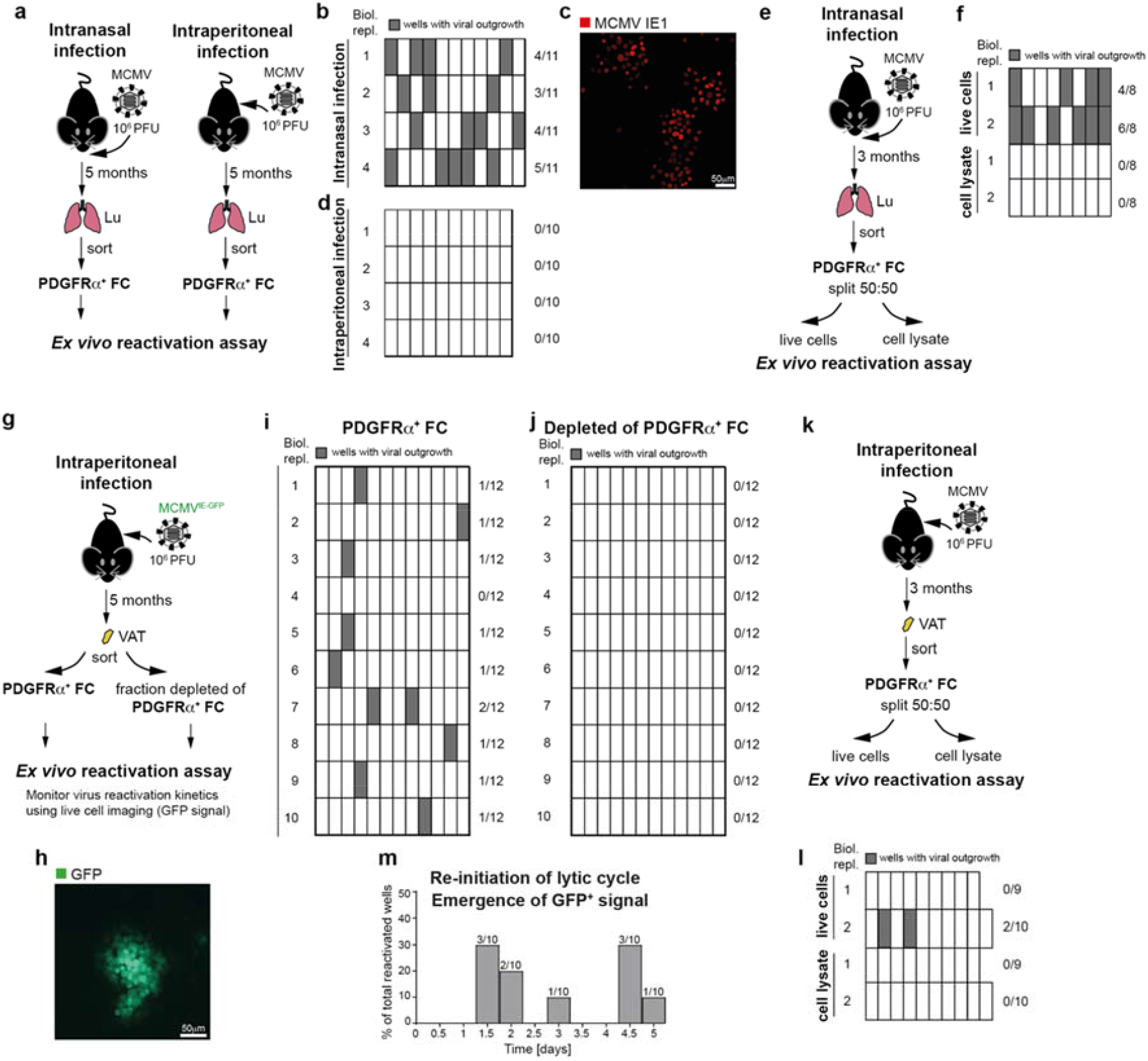
MCMV establishes latent infection in PDGFRα^+^ FC *in vivo*. *Ex vivo* virus reactivation assay from PDGFRα^+^ FC purified from **a-f**, the lungs of mice latently infected with MCMV administered intranasally or **g-m**, from the VAT of mice latently infected with MCMV administered intraperitoneally. **a**, Experimental set up for b-d. **e**, Experimental set up for f. **g**, Experimental set up for h-j, m. **k**, Experimental set up for l. **b, d, f, i, j, l**, Frequency of viral outgrowth from wells with indicated cells. Biological replicates are from 1-2 mice per replicate. Numbers on the right side are the proportion of wells with virus outgrowth for the indicated biological replicate. **c** Representative immunofluorescence image showing detection of viral IE1 protein. **h** Representative fluorescence microscopy image of a GFP^+^ plaque that arose in the *ex vivo* reactivation assay. **m** Kinetic of re-initiation of viral lytic cycle. Shown is the proportion of wells in which GFP signal had first emerged at the indicated time-point.

### PDGFRα^+^ FC are a site of lytic virus replication during primary MCMV infection

The apparent latent persistence of MCMV in PDGFRα^+^ FC *in vivo* implied the question whether or not they could also support productive virus cycle during primary MCMV infection. To directly assess the permissiveness of PDGFRα^+^ FC for lytic MCMV infection, we purified PDGFRα^+^ FC from the lungs of naive mice and infected them *ex vivo* with recombinant MCMV^tFP-GRB^, which expresses major IE promoter-driven green fluorescent protein (GFP) and late gene promoter-driven red fluorescent protein (mCherry) (Fig. 5a). Analysis of viral replication kinetics during *in vitro* culture demonstrated that PDGFRα^+^ FC are permissive for lytic MCMV infection (Fig. 5b-c). To address this *in vivo*, we first investigated if lytic gene expression was induced in PDGFRα^+^ FC during the acute phase of infection with MCMV. Given that spleen becomes infected with MCMV within hours following intraperitoneal inoculation [22], we isolated PDGFRα^+^ FC from the spleens of mice 12 hours post intraperitoneal infection, and analysed the expression of viral genes *ie1/*3, *M38* and *M48(SCP)* by RT-qPCR. At 12 hours post infection, splenic PDGFRα^+^ FC expressed high levels of all three viral transcripts at levels at least 1,000-fold higher than in any tested subset of PDGFRα^+^ FC from latently infected mice (Table 2, see also Table 1). Therefore, splenic PDGFRα^+^ FC supported lytic viral transcription shortly after the inoculation of mice with MCMV.

**Table 2.** RT-qPCR analysis of the expression of MCMV genes in PDGFRα^+^ FC at 12 hours post infection. cDNA isolated from pooled cell preparations from 4 mice was analysed in 2 technical duplicates per MCMV gene.

|  |  | Est. cell no. | MCMV transcript copies detected per reaction |  |  |  |  |  |
| --- | --- | --- | --- | --- | --- | --- | --- | --- |
|  |  |  | *ie1/3* |  | *M38* |  | *M48(SCP)* |  |
| 12 hours after intraperitoneal infection | Spleen PDGFRα<sup>+</sup> FC | 7,944 | 219,224 | 211,938 | 411,485 | 411,485 | 61,694 | 61,289 |
| uninfected | Spleen PDGFRα<sup>+</sup> FC | 6,113 | 0 | 0 | 0 | 0 | 0 | 0 |

**Figure 5.**
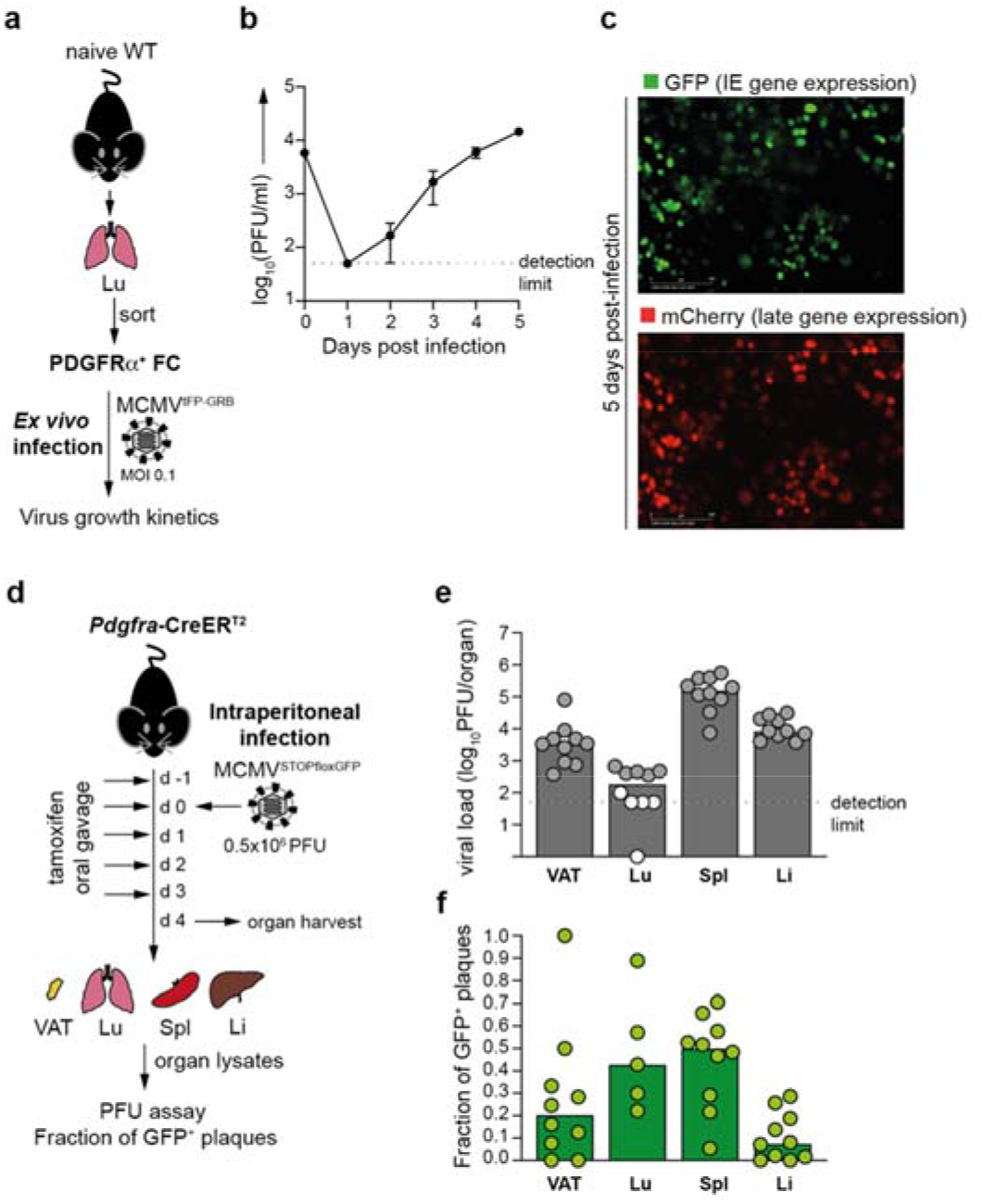
PDGFRα^+^ FC are a site of lytic virus replication during primary MCMV infection. **a-c**, Analysis of virus replication kinetics in lung-derived PDGFRα^+^ FC following their *ex vivo* infection of with MCMV^tFP-GRB^, which expresses IE promoter driven-green fluorescent protein (GFP) and late gene promoter-driven red fluorescent protein (mCherry). **a** Experimental set up for b-c. **b** Data are presented as arithmetic mean ± SD of n = 3 technical replicates and are from one representative experiment of 2 using cells sorted from pooled lung preparations from 3 mice. **c** Representative fluorescence microscopy images of GFP and mCherry expression 5 days after *ex vivo* infection. **d-f**, Contribution of PDGFRα^+^ FC to MCMV propagation *in vivo* determined after intraperitoneal infection of *Pdgfra*-CreER^T2^ (+/−) mice with a Cre-inducible reporter virus, MCMV^floxSTOP-GFP^. **d** Experimental set up for e-f. **e** Median virus titer in indicated organs. Circles represent samples from individual mice. Grey-filled circles are samples for which percentage of GFP^+^ plaques could be determined. White-filled circles are samples for which percentage of GFP^+^ plaques could not be determined. **f** Median proportion of GFP^+^ plaques in indicated organs. Circles represent samples from individual mice.

To confirm that PDGFRα^+^ FC also supported productive replication of the virus *in vivo*, we used mice heterozygous for *Pdgfra*-CreER^T2^ knock-in allele [40] (hereafter termed *Pdgfra*-CreER^T2^ (+/−), that express tamoxifen-activatable Cre recombinase from one endogenous *Pdgfra* locus while the other locus is preserved, and infected them with a reporter virus MCMV^floxSTOP-GFP^ [41]. MCMV^floxSTOP-GFP^ undergoes deletion of the stop cassette preventing GFP expression in cells that express active Cre recombinase and was used previously with Tie2-Cre or Alb-Cre mice to define virus replication in endothelial cells or in hepatocytes [11]. Following the infection of *Pdgfra*-CreER^T2^ (+/−) mice with MCMV^floxSTOP-GFP^ it is expected that the viruses that replicate in PDGFRα^+^ FC become genetically recombined so their progeny gives rise to GFP^+^ plaques on monolayers of MEFs. *Pdgfra*-CreER^T2^ (+/−) mice were infected with MCMV^floxSTOP-GFP^ via the intraperitoneal route on day 0, fed tamoxifen from −1 to 3 days post infection to activate the Cre recombinase, and sacrificed on day 4 post infection (Fig. 5d). Organ lysates from the spleen, VAT, liver and lungs were used to determine the frequency of GFP^+^ plaques. On day 4 post infection, GFP^+^ virus was detected as ~10% of plaques in the liver, ~20 % in the VAT and ~50 % in the spleen and lungs, although in the latter organ the percentage of GFP^+^ plagues could only be reliably measured in 5 out of 10 samples due to generally low infectious virus titer in this organ (Fig. 5e-f). The emergence of GFP^+^ virus was entirely dependent on the presence of Cre recombinase since similar virus titer but no GFP^+^ plaques were found in organs of infected Cre-negative littermate controls (data not shown).

Thus, multiple organs contained the progeny of viruses that had replicated in PDGFRα^+^ cells, indicating that PDGFRα^+^ FC play a role in MCMV *in vivo* replication. Of note, given that MCMV may have spread from the sites of initial replication within 4 days, it is not possible to distinguish whether the virus had replicated in PDGFRα^+^ FC present in the given organ or elsewhere. In sum, PDGFRα^+^ FC supported both productive infection and latent MCMV persistence *in vivo*.

### *Stat1* is required for MCMV genome persistence in fibroblastic cells *in vivo*

Next, we sought to identify host pathway/-s that affect MCMV’s ability to latently persist in PDGFRα^+^ FC *in vivo*. We hypothesized that the same signal/-s that inhibit lytic cycle entry by MCMV may concomitantly enable the virus to establish and/or maintain latent infection. Since IFNβ was sufficient to silence IE transcription [20], we chose to assess the putative involvement of STAT1, which crucially mediates responses to IFNs [16] by testing lytic and latent MCMV infection in *Stat1*^−/−^ mice. To circumvent the caveats linked to the increased sensitivity of *Stat1*^−/−^ mice to MCMV, we generated and used a single-cycle GFP-reporter MCMV mutant, which we name MCMV^ΔgLGFP^. To this effect we disrupted the gene for the essential virion glycoprotein L (gL/M115) in our GFP reporter virus, MCMV^IE1/3-GFP^ [29] on a Δm157 background. MCMVs lacking gL can be propagated in gL-complementing cells, to generate virus stocks consisting of pseudotyped gL^+^ virions that are able to infect and replicate for one cycle in gL deficient cells, whereupon their progeny virions will lack the gL and thus cannot spread further to other cells [42–44]. As shown in Fig. 6a, WT and *Stat1*^−/−^ mice were infected with MCMV^ΔgLGFP^, followed by analysis of lytic infection and latent persistence of the virus respectively at 22 hours and 60 days post infection in fibroblastic cells in the spleen, which are known to be directly infected by MCMV virions injected intraperitoneally or intravenously [22]. We focused specifically on red pulp FC, firstly because their high MCMV genome content during *in vivo* latency offered a broad dynamic range and secondly, to eliminate confounding effects due to a potential change in the composition of spleen stromal cells in *Stat1*^−/−^ mice. Since Ly6C is an IFN-regulated gene, Ly6C^+^ red pulp FC were identified as PDGFRα^+^BST-1^−^CD146^−^ FC, as previously reported [29]. The abundance of red pulp FC was comparable between WT and *Stat1*^−/−^ mice (Supplementary Fig. 2c). At 22 hours after intraperitoneal or intravenous injection of MCMV^ΔgLGFP^, *Stat1*^−/−^ mice showed a significantly higher percentage of GFP^+^ red pulp FC (Fig. 6b-d). Consistent with being lytically infected, GFP^+^ red pulp FC from WT and *Stat1*^−/−^ mice expressed high levels of *ie1/ie3, M38* and *M48(SCP)* transcripts (Table 3). Therefore, more red pulp FC became lytically infected in the absence of STAT1. At 60 days post infection, red pulp FC were devoid of GFP^+^ cells (Fig. 6f) and viral gene expression was silenced (Table 3). Strikingly, whereas MCMV genomes persisted in red pulp FC from WT mice at ~ 300 copies per 100,000 cells, these cells were profoundly depleted of MCMV genomes and contained only ~ 10 copies per 100,000 cells in *Stat1*^−/−^ mice (Fig. 6g). Together, the above data implicate a dual role of STAT1 as a negative regulator of lytic MCMV infection and a positive regulator of latent MCMV persistence in fibroblastic cells *in vivo*.

**Table 3.**
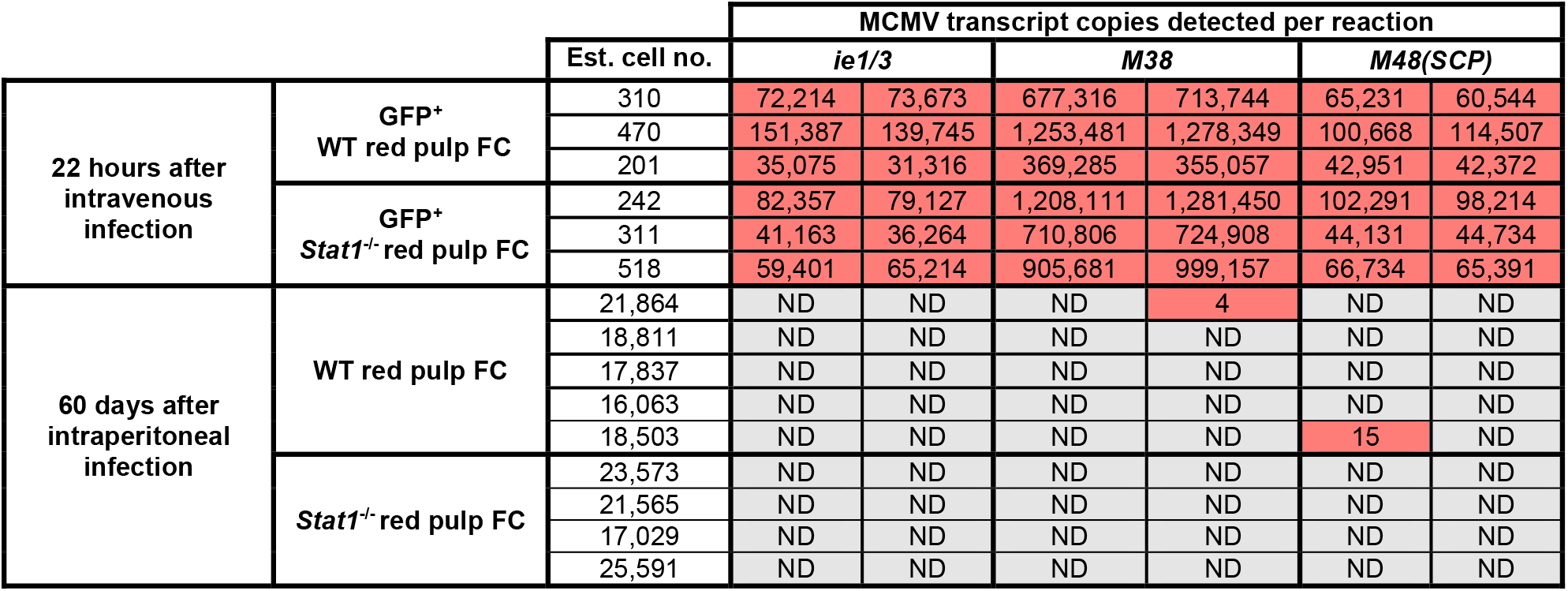
RT-qPCR analysis of the expression of MCMV genes in the indicated subsets purified from WT or *Stat1*^−/−^ mice that had been infected with MCMV^ΔgL-GFP^ for 22 hours or 60 days. Each row represents cDNA from a different biological replicate analysed in 2 reactions (technical duplicates) per MCMV gene. Each biological replicate represents cells sorted from a single mouse in a single experiment (22 hours post infection) or cells sorted from pooled preparations of 2 mice and are from 2 independent experiments (60 days post infection).

**Figure 6.**
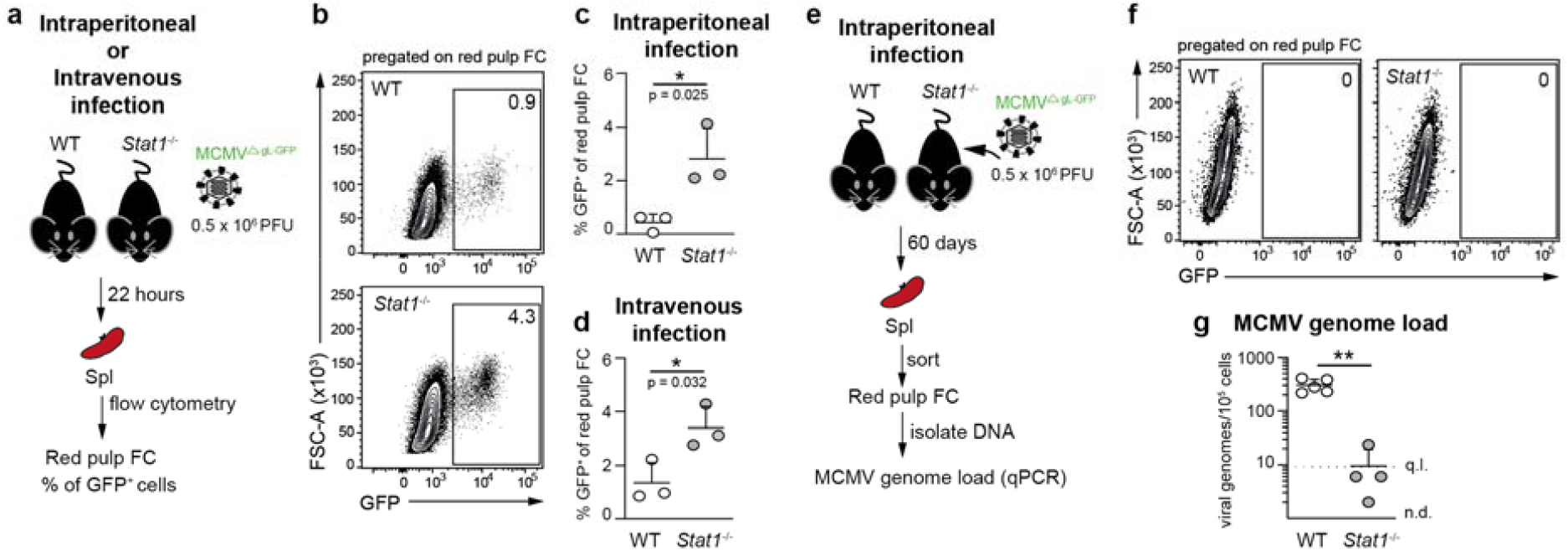
*Stat1* is required for MCMV genome persistence in fibroblastic cells *in vivo*. **a-d**, Analysis of the lytic infection of red pulp FC in WT and *Stat1*^−/−^ mice 22 hours after the administration of MCMV^ΔgL-GFP^ intraperitoneally or intravenously. **a** Experimental set up for b-d. **b-d** Percentage of GFP^+^ cells in red pulp FC. **b** Representative flow cytometry plot. Numbers indicate the percentage of gated cells. **c, d** Data are presented as the arithmetic mean ± SD of n = 3 biological replicates (individual mice) and are from 1 experiment performed for each infection route. *, p-value < 0.05 (exact p value indicated in the plot), unpaired t test. **e-g**, Analysis of latent MCMV persistence in red pulp FC from WT and *Stat1*^−/−^ mice 60 days after intraperitoneal administration of MCMV^ΔgL-GFP^. **e** Experimental set up for f-g. **f** Plots show the lack of GFP^+^ cells in red pulp FC. **g** Number of virus genomes per 100,000 of red pulp FC. Horizontal line depicts arithmetic mean ± SD from n = 4 biological replicates (depicted as symbols) and are pooled from 2 independent experiments. Biological replicates represent cells sorted from pooled preparations from 2 mice. **, p-value < 0.01, Welch’s t test. q.l., quantification limit; n.d., not detected.

## Discussion

This work provides to our knowledge the first comprehensive characterisation of the distribution of MCMV genomes in multiple tissue-resident cell types across numerous organs during MCMV latency *in vivo*, and the first demonstration of latent MCMV persistence in fibroblastic cells. Latent MCMV persistence in PDGFRα^+^ FC was not restricted to a single organ but represented a wide-spread phenomenon engaging many organs, suggesting that fibroblasts are a physiologically relevant, and in some organs may be a major site of MCMV *in vivo* latency. Our findings are not in opposition with previous identification of endothelial cells and macrophages as cellular sites of MCMV latency *in vivo* but rather confirm and extend previous knowledge by broadening the scope of tissues in which these cell types were tested for MCMV genome presence. In our model, which involved infection of adult C57BL/6 mice with tissue culture derived MCMV, we found the content of MCMV genomes in monocytes to be below 10 copies per 100,000 cells at 5 months post infection. This is similar to a recent study by Liu *et al*., in which 3 week-old BalbC mice were infected with MCMV Smith strain and analysed 3-6 months later, where the MCMV genome load in monocytes was more robust but still not higher than 10 copies per 100,000 cells [7]. It has been suggested that the presence of latent HCMV in endothelial cells may be a consequence of transdifferentiating latently infected myeloid progenitor cells [45], which might in theory explain our observations. However, the scarcity of viral genomes in monocytes in relation to tissue-resident cells, argues against this explanation in MCMV latency. Furthermore, given modern knowledge about the origin of fibroblastic cells in the mouse we deem this scenario very unlikely for MCMV. It is currently well established that CD45^−^PDGFRα^+^ or PDGFRβ^+^ fibroblastic cells found in mouse organs under homeostatic conditions are mesenchymal-derived and not a progeny of circulating progenitors originating from hematopoietic or mesenchymal BM-derived stem cells [27]. Although differentiation of monocyte/Mϕ-like cells into fibroblasts has been documented in the context of organ injury or fibrosis [46, 47], the low MCMV content of monocytes and Mϕ in most studied organs is at odds with the high enrichment of CD45^−^PDGFRα^+^ for MCMV genomes observed in our study. An alternative hematopoietic source of fibroblast-like cells in a perturbed state can be CD34^+^Col1a1^+^ fibrocytes, but they and their progeny were excluded from our analysis based on their positivity for CD45 [48]. Thus, available indirect evidence strongly suggests that MCMV not only latently persist in but also latently infects CD45^−^PDGFRα^+^ FC.

Notably, CD45^−^PDGFRα^+^ FC also became lytically infected by the virus and contributed to its *in vivo* replication during the acute stage following primary infection. These findings shed new light on the role of fibroblasts in MCMV’s life cycle *in vivo* by showing that not only does this cell type support viral lytic cycle, as previously understood, but it nevertheless serves as the site of latent MCMV persistence. It is important to note that the above conclusion can only be inferred for the entire population of PDGFRα^+^ FC from a given organ, but not on a single-cell basis. PDGFRα^+^ FC are known to be heterogenous [32] and it is entirely possible that lytic and latent infection are allocated to distinct subsets or maturation stages. Additional studies involving single cell transcriptomics will be required to resolve this matter in the future.

While our study was focused on demonstrating bone-fide latent virus persistence in fibroblastic cells and identification of a cellular signalling pathway on which it is critically dependent, some important follow up questions remain that lie outside of the scope of the current project. For instance, although our mechanistic studies in *Stat1*^−/−^ mice unravelled that endogenous *Stat1* was required to enable latent persistence of MCMV in fibroblastic cells, we did not determine whether or not the requirement for *Stat1* was cell intrinsic. To decipher this, mice bearing conditional deletion of *Stat1* in fibroblasts will need to be implemented. Should *Stat1* be specifically required in fibroblasts, it will further need to be clarified whether or not *Stat1* affects the ability of the virus to establish latent infection or rather persist in a latent form, for example by preventing spontaneous reactivation.

Addressing these questions will likely require careful probing of single-cell transcriptomes of fibroblastic cells at various time points after infection *in vitro* or *in vivo*. Regardless of the results of the future investigations addressing these outstanding questions, it is important to observe that the data generated in the course of this study unravel that one of the most prominent antiviral host factors, which is crucial for the resolution of virus infections and host survival, concomitantly enables a herpesvirus to latently persist in its host, at least in certain cell types. In sum, the findings of this study significantly broaden our understanding of the functions of murine *Stat1* during *in vivo* infection with a β-herpesvirus.

## Methods

### Mice

Experiments were performed with female mice on C57BL/6 background. Mice were infected at 8-10 weeks of age, except in Fig. 6 where the infection was done at 14 weeks of age. C57BL/6JrJ mice (purchased from Janvier Labs), *Pdgfra*-CreER^T2^ mice (IMSR_JAX:032770) [40] and *Stat1*^−/−^ mice (generated by mating *Stat1*^flox^ mice with CMV-Cre deleter strain) [49] were bred and maintained under specific pathogen free (SPF) conditions according to Federation of European Laboratory Animal Science Associations (FELASA) guidelines respectively at central animal facility of HZI Braunschweig, Germany; Faculty of Medicine, University of Rijeka, Croatia, and University of Veterinary Medicine Vienna, Austria. Animal procedures were approved as due by The Lower Saxony State Office of Consumer Protection and Food Safety; by The Ethics Committee at the Faculty of Medicine, Rijeka and Ethics Committee of the Veterinary Department of the Ministry of Agriculture, Croatia, and by the Ethics and Animal Welfare Committee of the University of Veterinary Medicine Vienna and the Austrian Federal Ministry of Science and Research. Virus was administered in 200 µl PBS intraperitoneally or in 20 μl in PBS intranasally as described previously [50]. *Pdgfra*-CreER^T2^ mice received tamoxifen (0.1mg/g; Sigma) diluted in corn oil by oral gavage daily from day −1 to day 3 post infection.

### Viruses

Viral stocks were generated in tissue culture by transfecting MCMV genome-carrying BACs into NIH3T3 cells (ATCC CRL-1658) using FuGene HD (Promega). Reconstituted viral particles were passaged five times on M2-10B4 cells (ATCC CRL-1972) before generating sucrose cushion-purified viral stocks. All MCMVs used in this study were based on the Mck2-repaired BAC-encoded MCMV strain, clone pSM3fr-MCK-2fl 3.3 [51]. MCMV^IE-GFP^ [29] and MCMV^floxSTOP-GFP^ [41] were described before. MCMV^tFP-GRB^ expressing three different fluorescent proteins under the control of endogenous viral promoters was generated by *en passant* mutagenesis [52] using expression cassettes described earlier [53]. In short, constructs encoding fluorescent reporters (EGFP, mTagBFP-NLS and mCherry) plus a distal P2A-encoding sequence were inserted into the start codon of IE1/3 (EGFP), E2 (mTagBFP-NLS), and SCP (mCherry). MCMV^ΔgLGFP^ was generated by disrupting m157 and gL/M115 genes in MCMV^IE-GFP^. Viral genes were disrupted by frameshift mutations induced by replacement of 38 bp of each gene’s open reading frame (ORF) with two STOP codons targeted to the ORF’s start (gL) or after codon 22 (m157). MCMV^ΔgLGFP^ was propagated in gL-expressing NIH-3T3 cell line [42] (a kind gift from Ian Humphreys, Cardiff University).

### Cell Isolation

Stromal-vascular fractions from VAT were prepared as previously described [35] except that liberase TM in the digestion medium was replaced with collagenase P (0.4 mg/ml). Spleen, LNs and SGs were processed according to a previously established protocol [29]. In short, enzymatic treatment with collagenase P (0.4 mg/ml), dispase II (2 mg/ml) and DNase I (50 µg/ml) was performed for 30 min at 37 °C and then repeated for additional 20 min, followed by incubation with 5 mM EDTA for 5 min. CD45^−^ cell fractions from the spleen and LNs were enriched by depletion of CD45^+^ cells while splenic Mϕ were enriched by positive selection of VCAM-1^+^ cells using MACS (Miltenyi Biotech). Lungs were injected with 2.5 ml digestion solution containing collagenase P (0.4 mg/ml), dispase II (2 mg/ml) and DNase I (50 µg/ml), minced with scissors and then processed in gentleMACS Dissociator (Miltenyi Biotech) using program “37C_m_LDK_1”. Liver digestion was performed using the retrograde perfusion technique via the inferior vena cava described by Mederacke *et al*. [30] with modifications. Briefly, organs were *in situ* perfused with 10 ml of pre-warmed Liver Perfusion Medium (Gibco) followed by injection of 20 ml of pre-warmed digestion medium (1 mg/ml collagenase P, 1 mg/ml dispase II, 50 µg/ml DNase I in Gibco Hank’s Balanced Salt Solution (HBSS) with 10 mM HEPES and 1.5 mM calcium chloride) over 5-7 min. Afterwards, organs were dissected, minced with scissors and *in vitro* digested for 45 min at 37°C. Non-parenchymal cells were enriched by two rounds of centrifugation at 50 × g for 5 min whereupon only supernatant was collected.

### Flow Cytometry and Cell Sorting

Antibodies used in this study are listed in Supplementary Table 1. Dead cells (identified using 7-AAD or Zombie Fixable Viability Kits; both from BioLegend) and cell aggregates (identified on FSC-A versus FSC-H scatter plots) were excluded from all analyses. Sorting was performed on ARIA-Fusion, Aria-II SORP or Aria-II (BD Biosciences). Analysis was done using FlowJo software (version 10.8.1, BD Biosciences).

### Immunofluorescence

Cells were fixed in 4 % (wt/vol) paraformaldehyde, permeabilized in 0.1% Triton-X and blocked with 2% BSA. Staining with anti-IE1 antibody (IE 1.01, CapRi) was done overnight at 4°C and was followed by incubation with anti-mouse IgG (H+L) F(ab’)_2_ conjugated to Alexa Fluor 647 (Cell Signaling Technology). Images were acquired with ZEISS LSM 980 laser-scanning confocal microscope and analysed using ZEN software (Carl Zeiss MicroImaging).

### Quantification of MCMV Genome Load by qPCR

Quantification of MCMV genome load was performed according to the method described by Simon *et al*. [33] with modifications. Briefly, DNA was extracted using AllPrep DNA/RNA Micro Kit (QIAGEN), and after elution precipitated with ethanol, washed, and resuspended in 40 µl dH_2_O. Samples were assayed for viral gB and mouse Pthrp DNA copies in technical duplicates using 9 µl of sample DNA per reaction. Serial dilutions of 10^6^-10^1^ copies per reaction of the pDrive_gB_PTHrP_Tdy plasmid [33] were used to generate standard curves for both qPCR reactions. Dilutions of 10^4^-10^1^ plasmid copies contained 1 ng carrier RNA (QIAGEN). QPCR was performed in a LightCycler 480 Instrument II (Roche) using Fast Plus EvaGreen qPCR master mix (Biotium). Cycling conditions were as follows: enzyme activation, 2 min 95°C, followed by 50 cycles of denaturation for 10 sec at 95°C, annealing for 20 sec at 56°C and extension for 30 sec at 72°C. Specificity of amplified sequences was validated by inspection of melting curves. Primer sequences: *Pthrp* 5’-ggtatctgccctcatcgtctg-3’ and 5’-cgtttcttcctccaccatctg-3’; gB 5’-gcagtctagtcgctttctgc-3’ and 5’-aaggcgtggactagcgataa-3’.

### Analysis of MCMV Transcripts by RT-qPCR

Total RNA was extracted using AllPrep DNA/RNA Micro Kit with on-column DNA removal (QIAGEN). cDNA was synthesized with SuperScript IV Reverse Transcriptase and a 1:1 mixture of oligo-dT and random oligonucleotide hexamers (Invitrogen). QPCR was performed in a LightCycler 480 Instrument II (Roche) using Fast Plus EvaGreen qPCR master mix (Biotium). Serial dilutions of 10^6^-10^1^ copies per reaction of MCMV genome containing BAC (clone pSM3fr-MCK-2fl 3.3 [51]) were used to generate standard curves. Cycling conditions were as follows: enzyme activation, 2 min 95°C, followed by 50 cycles of denaturation for 10 sec at 95°C, annealing for 20 sec at 57°C and extension for 30 sec at 72°C. Specificity of amplified sequences was validated by inspection of melting curves. RT-qPCR reactions prepared from equal amount of sample RNA but with the omission of reverse transcriptase served as controls for the absence of carry-over viral DNA. Primer sequences: *Gapdh* 5’-tgtgtccgtcgtggatctga-3’ and 5’-cctgcttcaccaccttcttgat-3’; ie1/3 5’-gggcatgaagtgtgggtaca-3’ and 5’-cgcatcgaaagacaacgcaa-3’; M38 5’-cccatgtccacgttttggtg-3’ and 5’-tctcaagtgcaatcgcctca-3’; M48/SCP 5’-aggacgtgttgaacttcttctcc-3’ and 5’-tcgatgtccgtctccttcac-3’.

### *Ex vivo* Reactivation Assay

FC were seeded into fibronectin-coated 96 Half Area Well Flat Bottom Microplates (Corning) (lung FC) or gelatin- or fibronectin-coated 96 Well Flat Bottom Microplates (Greiner Bio-One) (VAT FC) at a density of 100,000 cells per well in culture medium (high-glucose DMEM supplemented with 1 % (vol/vol) GlutaMAX, 10 mM HEPES, 50 µM 2-mercaptoethanol, 200 U/ml penicillin, 200 U/ml streptomycin, 50 µg/ml gentamicin and 5 % FCS) and incubated in a humidified incubator under 5 % CO_2_ and at 37°C for 7 days without replacing the medium. Cell lysates prepared in gentleMACS M Tubes for tissue homogenisation using gentleMACS Dissociator (Miltenyi Biotech, program “mlung_02”) were assayed in parallel by incubation with MEFs. To track the emergence of GFP^+^ cells, plates were imaged with IncuCyte S3 automated fluorescence microscope (Sartorius). After 7 days, plates were directly inspected for the presence of plaques and additionally, the culture supernatant from each well was tested for the presence of infectious virus by plaque assay on MEFs. To ensure detection of even minimal amounts of virus [54], following addition of the supernatant, MEFs were spun down at 800 g for 30 min and then incubated at 37°C for another 30 min. Afterwards, the inoculum was removed and the cells overlaid with medium containing 0.75 % (wt/vol) methylcellulose (Sigma). After 4 days at 37 °C, the plates were examined for the presence of plaques under an inverted microscope.

### *Ex vivo* Infection of Cells

Lung-derived FC at 80 % confluency were infected with MCMV^tFP-GRB^ at an MOI of 0.1 for an hour at 37°C. Afterwards, the medium was changed to cultivation medium (high-glucose DMEM (Gibco) supplemented with 1 % (vol/vol) GlutaMAX, 10 mM HEPES, 50 µM 2-mercaptoethanol, 200 U/ml penicillin, 200 U/ml streptomycin, 50 µg/ml gentamicin and 5 % FCS). Supernatants were collected from a different set of wells every 24 hours until 5 days post infection and stored at −80°C. Titration of the virus was done by infecting MEF cell monolayers grown in 48-well plates with serially diluted supernatants for an hour at 37 °C. Thereupon, the inoculum was removed, and the cells were overlaid with culture medium containing 0.75 % (wt/vol) methylcellulose (Sigma). Plaques were counted following incubation of cells for 4 days at 37 °C.

### Statistics and Reproducibility

Unpaired t test, Welch’s t test and one-way ANOVA with Dunnett’s multiple comparisons test were performed with GraphPad Prism 9. No values were excluded from statistical analysis as outliers. P-values < 0.05 were considered significant. * p<0.05; ** p<0.01; *** p<0.001; **** p<0.0001. Error bars denote mean ± SD. Sample size is reported in figure captions. Results were reproduced by two or more independent experiments performed.

## Data Availability

Source data are provided with this paper.

## Acknowledgements

This study was funded by the Deutsche Forschungsgemeinschaft (DFG, German Research Foundation) – Projektnummer 158989968 - SFB 900; the RESIST Excellence cluster (EXC 2155, project B6); the German Centre for Infection Research (DZIF) of the German Federal Ministry of Science and Education (BMBF) through TTU 07.834 (to L.C-S.) and the European Union’s Horizon 2020 research and innovation program under Grant Agreement 793858 (to K.M.S.). Ilija Brizić is supported by Research Cooperability program of the Croatian Science Foundation funded by the European Union from the European Social Fund under the Operational Programme Efficient Human Resources 2014–2020 (PZS-2019-02-7879). We thank Inge Hollatz-Rangosch and Ayse Barut for technical assistance. We acknowledge Lothar Gröbe and Maria Höxter from HZI Core Flow Cytometry Facility as well as Marina Pils, Katrin Schlarman and Petra Beyer from HZI Animal Facility. We thank Mathias Müller and Birgit Strobl (University of Veterinary Medicine Vienna) for providing *Stat1*^−/−^ mice. We acknowledge Caroline Lassnig for providing help with *in vivo* experiments. We thank Hansjörg Hauser for useful discussions.

## Author contributions

Concept and study design, K.M.S. and L.C-S.; Investigation, K.M.S., F.K., N.G., H.M., Y.K.; Methodology, K.M.S. and L.C-S., Materials: U.R. and T.K.; Resources, L.C-S. and I.B.; Funding acquisition, L.C-S., K.M.S. and I.B.; Writing of the manuscript, K.M.S. and L.C-S.; Supervision, K.M.S. and L.C-S.

