## supplementary data for "Fibroblastic cells are a site of mouse cytomegalovirus *in vivo* lytic replication and latent persistence oppositely regulated by *Stat1*"

Supplementary Figure 1

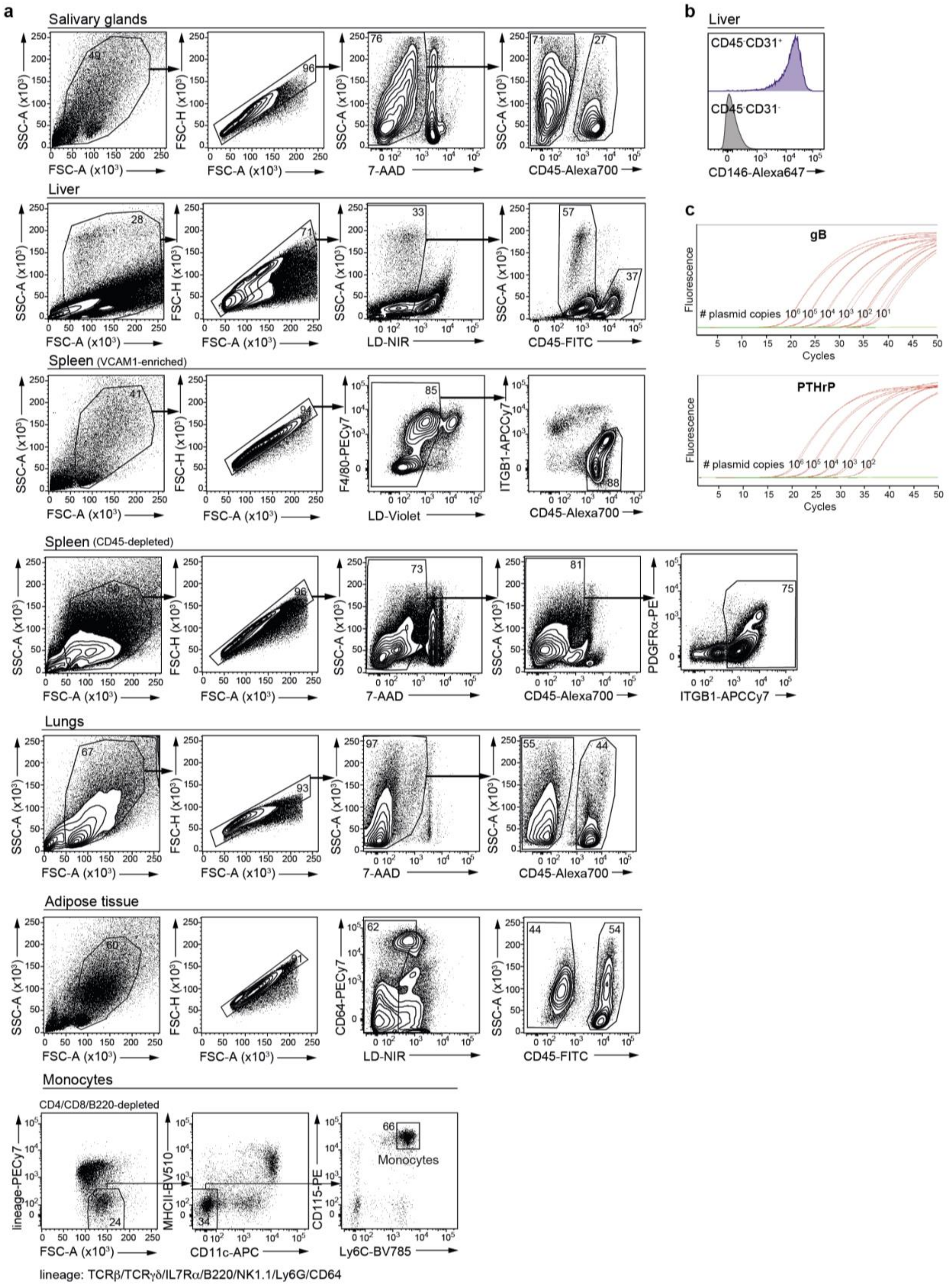

**Supplementary Figure 1**
**a** Pre-gating strategy used for sorting of the indicated cell subsets. **b** CD146 expression by liver EC. Representative plots from two mice. **c** Dynamic range of the qPCR assay for quantification of MCMV and mouse genome copies validated using serial dilutions of a plasmid with inserted viral gB and mouse *Pthrp* genomic sequences.

**Supplementary Figure 2**

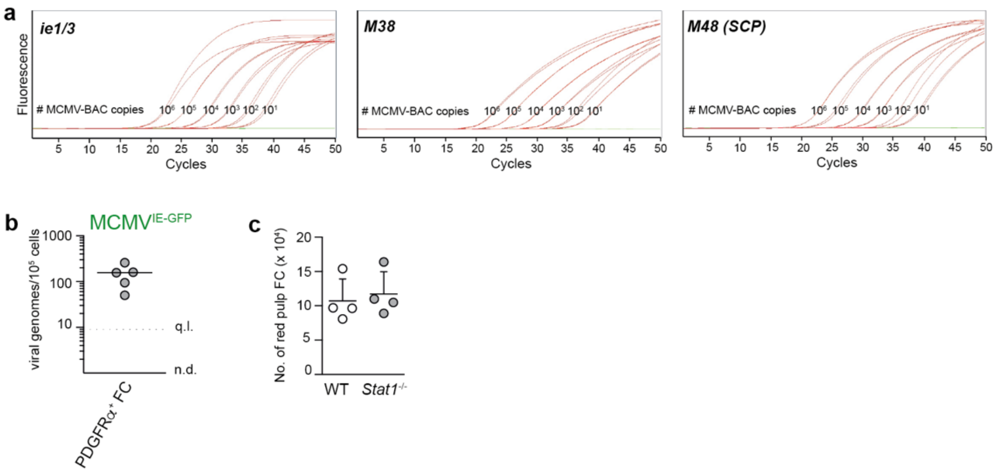


**Supplementary Figure 2**

**a** Dynamic range of the qPCR assay for quantification of ie1/3, M38 and M48 cDNA copies validated using serial dilutions of MCMV BAC. **b** MCMV genome load in PDGFRα$^+$ FC purified from the AT of mice latently infected with $10^6$ PFU of MCMV$^{IE\text{-}GFP}$ administered intraperitoneally 5 months prior. Horizontal line depicts median from n = 5 mice (depicted as symbols). q.l., quantification limit; n.d., not detected. **c** Number of red pulp FC in 14 weeks old WT or *Stat1*$^{-/-}$ mice. Data are presented as the arithmetic mean ± SD of n = 4 biological replicates (individual mice) and are from a singular experiment.

### Supplementary Table 1. Antibodies

| Antibody | Source | Cat. No. |
|---|---|---|
| APC/Cy7 anti-mouse ITGB1 (clone HMβ1-1) | BioLegend | Cat# 102226 |
| Alexa Fluor 700 anti-mouse CD45.2 (clone 104) | BioLegend | Cat# 109822 |
| FITC anti-mouse CD45.2 (clone 104) | BioLegend | Cat# 109806 |
| FITC anti-mouse BST1 (clone KT157) | eBioscience | Cat# MA5-17948 |
| APC anti-mouse BST1 (clone BP-3) | BioLegend | Cat# 140208 |
| PE anti-mouse PDGFRα (clone APA5) | BioLegend | Cat# 135906 |
| APC anti-mouse PDGFRα (clone APA5) | BioLegend | Cat# 135908 |
| PE anti-mouse PDGFRβ (clone APB5) | BioLegend | Cat# 136006 |
| PE/Cy7 anti-mouse CD146 (clone ME-9F1) | BioLegend | Cat# 134714 |
| Alexa Fluor 647 anti-mouse CD146 (clone ME-9F1) | BioLegend | Cat# 134717 |
| Brilliant Violet 510 anti-mouse Ly6C (clone HK1.4) | BioLegend | Cat# 128033 |
| PE/Cy7 anti-mouse Ly6C (clone HK1.4) | BioLegend | Cat# 128018 |
| BD Horizon BV421 anti-mouse CD31 (clone MEC13.3) | BD Horizon | Cat# 562939 |
| PE/Cy7 anti-mouse CD31 (clone MEC13.3) | BioLegend | Cat# 102524 |
| PE anti-mouse VCAM1 (clone 429 (MVCAM.A)) | BioLegend | Cat# 105714 |
| PE/Cy7 anti-mouse F4/80 (clone BM8) | BioLegend | Cat# 123113 |
| APC anti-mouse F4/80 (clone BM8) | BioLegend | Cat# 123115 |
| APC/Cy7 anti-mouse EpCAM (clone G8.8) | BioLegend | Cat# 118218 |
| PerCP/Cy5.5 anti-mouse Ly6G (clone 1A8) | BioLegend | Cat# 127616 |
| PerCP/Cy5.5 anti-mouse Ly6C (clone HK1.4) | BioLegend | Cat# 128012 |
| PerCP/Cy5.5 anti-mouse Siglec-F (clone S17007L) | BioLegend | Cat# 155526 |
| PE anti-mouse Siglec-F (clone S17007L) | BioLegend | Cat# 155505 |
| Brilliant Violet 510 anti-mouse/human CD11b (clone M1/70) | BioLegend | Cat# 101263 |
| PE/Cy7 anti-mouse CD64 (clone X54-5/7.1) | BioLegend | Cat# 139314 |
| Brilliant Violet 510 anti-mouse CD11c (clone N418) | BioLegend | Cat# 117353 |
| PE anti-mouse CD115 (clone AFS98) | BD Pharmingen | Cat# 566839 |
| Brilliant Violet 785 anti-mouse Ly6C (clone HK1.4) | BioLegend | Cat# 128041 |
| APC anti-mouse CD11c (clone N418) | BioLegend | Cat# 117309 |
| Brilliant Violet 510 anti-mouse I-A/I-E (clone M5/114.15.2) | BioLegend | Cat# 107636 |
| FITC anti-mouse/human CD11b (clone M1/70) | BioLegend | Cat# 101205 |
| PE/Cy7 anti-mouse TCR β chain (clone H57-597) | BioLegend | Cat# 109221 |
| PE/Cy7 anti-mouse TCR γ/δ (clone GL3) | BioLegend | Cat# 118123 |
| PE/Cy7 anti-mouse IL-7Rα (clone S18006K) | BioLegend | Cat# 158209 |
| PE/Cy7 anti-mouse/human CD45R/B220 (clone RA3-6B2) | BioLegend | Cat# 103221 |
| PE/Cy7 anti-mouse NK-1.1 (clone PK136) | BioLegend | Cat# 108713 |
| PE/Cy7 anti-mouse Ly6G (clone 1A8) | BioLegend | Cat# 127618 |
